# Coverage landscape of the human genome in nucleus DNA and cell-free DNA

**DOI:** 10.1101/2024.12.03.626615

**Authors:** Jiaqi Luo, Shuai Cheng Li

## Abstract

For long, genome-wide coverage has been used as a measure of sequencing quality and quantity, but the biology hidden beneath has not been fully exploited. Here we performed a comparative analysis on genome-wide coverage profiles between nucleus genome DNA (gDNA) samples from the 1000 Genomes Project (n=3,202) and cell-free DNA (cfDNA) samples from healthy controls (n=113) or cancer patients (n=362). Regardless of sample type, we observed an overall conserved landscape with segmentation of coverage, where adjacent windows of genome positions present similar coverage. Besides GC-content, we identified protein-coding gene density and nucleosome density as major factors influencing the coverage of gDNA and cfDNA, respectively. Differential coverage of cfDNA vs gDNA was found in immune-receptor loci, intergenic regions and non-coding genes, reflecting distinct genome activities in different cell types. A further rise in coverage at non-coding genes and intergenic regions plus a further drop of coverage at protein-coding genes and genic regions within cancer cfDNA samples indicated a loss of contribution by normal cells. Importantly, we observed the distinctive feature of coverage convergence in cancer-derived cfDNA, with the extent of convergence positively correlated to stages. Based on the findings, we developed and validated an outlier-detection approach for cfDNA-based cancer screening without the need of cancer samples for training, outperforming current benchmarks on condition-matched and condition-unmatched cancer detection tasks.

## Introduction

Cell-free DNA (cfDNA) is one of the most sought-after biomarkers for the detection and monitoring of cancers or other diseases. A single tube of peripheral blood can extract both peripheral blood mononuclear cells (PBMC) that carry reference nucleus genome DNA (gDNA) and plasma cfDNA that potentially contains tumour-derived DNA with cancer-related mutations (Figure 1). This paired mode of somatic mutation calling has become a mature strategy widely used in clinical settings for the identification of targetable mutations or the monitoring of cancer progression (1–3). However, only a part of patients, more likely in late stages, carry known driver mutations related to cancers, and not all mutations are detectable in cfDNA. The phenomenon of clonal hematopoiesis (CH) where somatic mutations that could be shed into cfDNA accumulate in hematopoietic stem cells, may introduce ‘false positive’ in known oncogenes such as TP53, KRAS, GNAS, NRAS and PIK3CA (4) as it also occurs in healthy individuals. To expand the scope of liquid biopsy based on cfDNA, alternatives are required to circumvent the limitations of somatic mutation calling. Intrinsic connections were discovered between nucleus gDNA and cfDNA, that cfDNA fragments of variable lengths are the products of nuclease activities on gDNA released by cell death (5, 6). The parts of the gDNA under protection of protein-binding (e.g. nucleosomes, transcription factors) remain intact, while the rest being naked are susceptible to nuclease cleavage, especially at DNase I hypersensitive sites (DHSs). Genomic regions enriched with long fragments (120-180bp) generally map the positions of nucleosomes while those with dense short fragments (35-80bp) correlate to DHSs and/or transcription factor binding sites. A hallmark of actively transcribed genes is the reduction of nucleosome occupancy at and nearby the promoter region, leading to a drop in read coverage. The nucleosome and DHS positioning differs between tissues or cell types, thus providing the possibility to infer the (tumour) tissue-of-origin. Besides, significant differences in fragment sizes, end points and end motifs have been found between tumour-derived cfDNA and normal (control) cfDNA. These fragmentomic features coupled with machine learning models yield good performances on single datasets where train and test samples come from the same study, despite the generalizability to independent external datasets were not fully evaluated (7–9).

**Figure 1.**
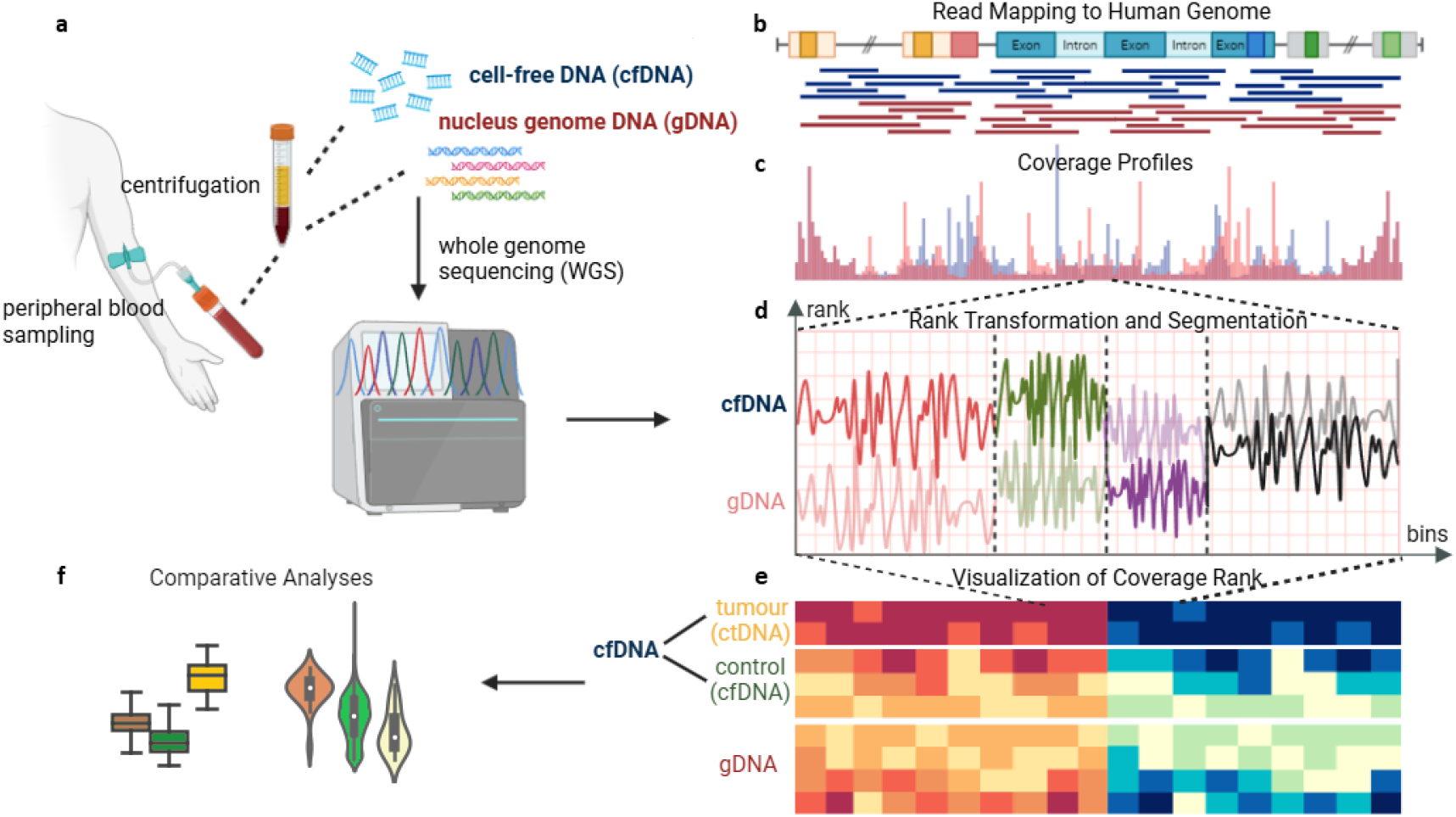
A graphical summary of the study design. **(a)** A single tube of peripheral blood can extract both nucleus genome DNA (gDNA) and cell-free DNA (cfDNA) for whole genome sequencing (WGS). We obtained WGS data of gDNA from 1000 Genomes (n=3,202), WGS data of cfDNA by cancer patients (n=362) and/or healthy controls (n=113) from the European Genome Achieve. **(b)** Raw data was mapped to the human reference genome hg38, and **(c)** coverage over 10kb windows were computed as genome-wide coverage profile. **(d)** Rank transformation was performed to transform coverage into rank to allow integration and comparison of data from different studies. We described the observation of rank segmentation, where adjacent windows share similar levels of coverage ranks. We identified borders of the rank segmentation, resulting in 1,090 coverage segments representing the genome-wide coverage landscape. **(e)** The coverage landscapes of gDNA and cfDNA of tumour (hereafter short-named as **ctDNA**) or control samples (hereafter short-named as **cfDNA**) were visualized with heatmaps, and were **(f)** systemically compared. A proof-of-concept method based on outlier-detection was developed and validated based on the distinctive feature identified in cancer samples.

Although used locally as an auxiliary variable to infer copy-number variation, protein-binding, transcriptions or tissue-of-origin in previous studies (1, 3, 5, 6), coverage itself has not been studied as an outcome variable at a genome-wide level. Coverage measures the number of sequencing reads that ‘covers’ a specified genomic position or region. Coverage information has been widely used as a quality control metric in sequencing experiments and mutation calling algorithms (10, 11). Lindner et al (12) introduced the use of genome coverage profiling to solve complex mixtures in metagenomics. The genome-wide coverage was read-based, regardless of fragment sizes. Inspired by previous work, we viewed the genome-wide coverage as an outcome of genomic, transcriptomic and epigenomic activities dictated by cell types, which may be biologically meaningful and distinguishable between health and diseases. We systemically compared and contrasted genome-wide coverage between gDNA versus cfDNA, and between cfDNA of healthy controls versus various cancers (Figure 1). We looked for coverage patterns that are conserved or distinct among sample types, and tried to explain the patterns with factors such as cell types, GC-content, positioning of genic/intergenic regions, protein-coding/non-coding genes and nucleosomes. Finally, we examined the feasibility of cfDNA-based cancer screening by genome-wide coverage profiling. Unlike copy-number analysis which relies on ploidy assumption, the coverage analysis in this study assumed real number ranges instead of integer multiples, offering potentially higher resolution on differentiating tumour and normal samples.

This study includes a lot of comparisons between cfDNA samples of cancer versus control. For the ease of expression, hereafter we use the term <u>***ctDNA***</u> to represent the <u>entire</u> cfDNA sample drawn from **tumour** patients, which differs from its conventional meaning; and <u>***cfDNA***</u>, if without specific clarification, to represent the <u>entire</u> cfDNA sample from healthy **controls**.

## Methods

### Datasets

Data used in this study included whole-genome sequencing (WGS) data of the 3,202 gDNA samples from the 1000 Genomes Project (1kGP) (13) and the ctDNA samples with or without condition-matched controls from multiple studies (9, 14–17) that have been deposited to the European Genome Archive (Table 1).

**Table 1.**
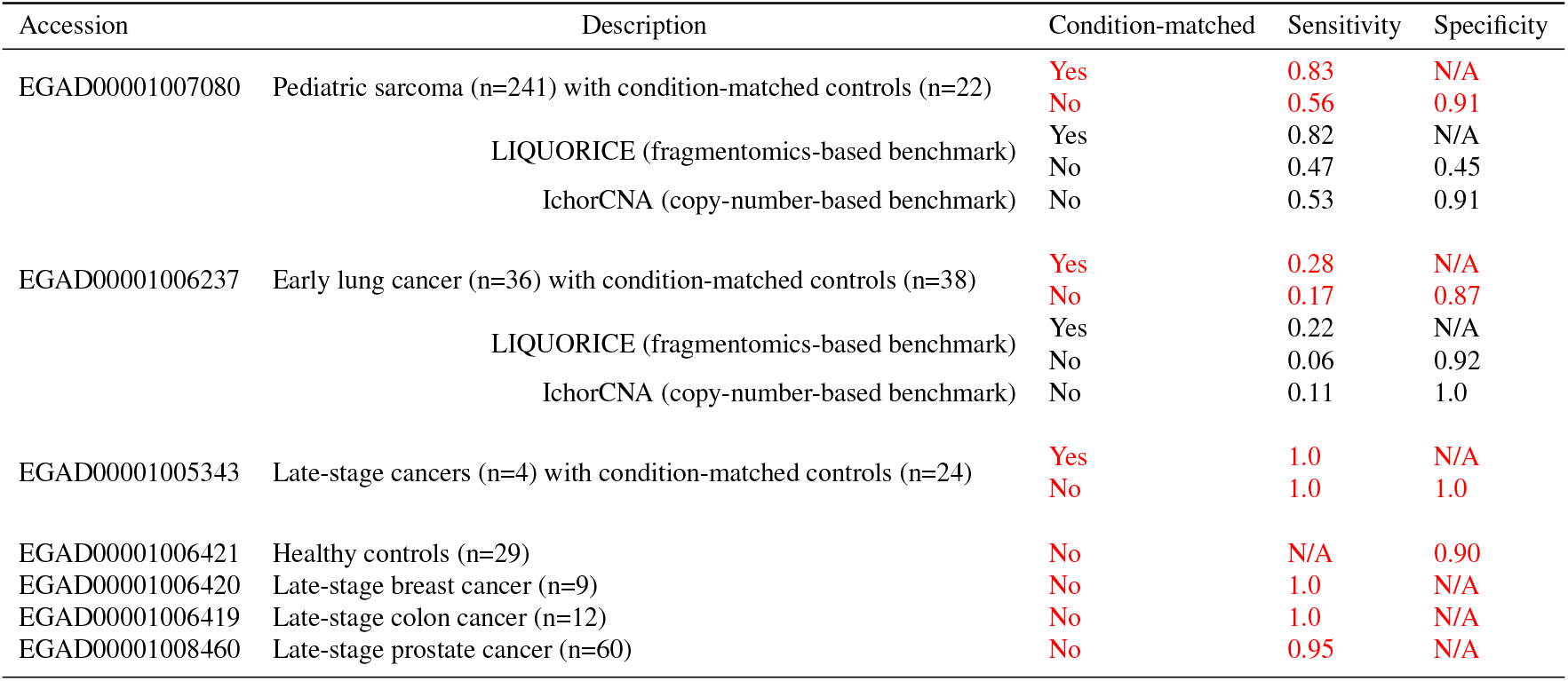
Performances of the proposed convergence-based outlier-detection method (highlighted in red) on various cancer-screening tasks.

### Data processing

The raw WGS data (fastq, bam) was pre-processed following the GATK best-practice pipeline (18). In brief, each sample was mapped to the hg38 human reference genome, with sequencing duplicates removed and base quality score recalibrated.

### Genome-wide coverage profiling

#### 10kb-window coverage fraction

Coverage at each genomic position was calculated with Samtools using the command ‘samtools depth -J -s’ (19). Coverage of all positions within each 10kb-window was sum together as the window’s coverage. Each window coverage was then divided by the total sum of all window coverage to get the coverage fraction. Note that repeated sequences located in pericentromeric regions can vary greatly between samples and affect the coverage fractions of the rest of the genome. Therefore, the three cytobands closest to the centromere from both arms of each chromosome were excluded from calculations. Non-autosomal chromosomes (chrX, chrY, chrM) were also excluded for coverage calculations to remove bias caused by gender. Finally, windows showing zero coverages in all samples were filtered, resulting in a total of 236,239 windows.

#### Rank transformation

A normalization/integration approach is required for the correct comparisons of samples within and between studies. Rank transformation was performed on the coverage fraction profile of each sample. The genome-wide, 10kb-windows of coverage fraction were ranked, the bigger the coverage fraction value, the bigger the rank; tied elements were assigned the possible minimum rank. Formally, for each window i with coverage fraction *f*_*i*_, the rank-transformed value *r*_*i*_ is formulated as:

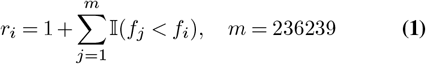

where 𝕀 is an indicator function:

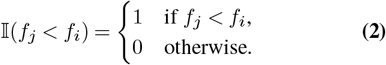

The rank transformation was compared with the raw coverage fraction, Z-score standardization and min-max normalization on dealing with batch effects. Among the methods compared, the rank transformation had been shown to minimize the differences between four datasets of control cfDNA samples (Supplementary Figure 1). To evaluate post-transform batch-effects, we performed principle component analysis on the rank-transformed data (Supplementary Figure 2), and inspected inter- and intra-dataset variations via heatmap for each chromosome (Supplementary Figure 3). From the heatmap we observed an overall inter-dataset consistency across the four control cfDNA datasets for each chromosome, except chr19, which exhibited significant inter- and intra-dataset variation.

**Figure 2.**
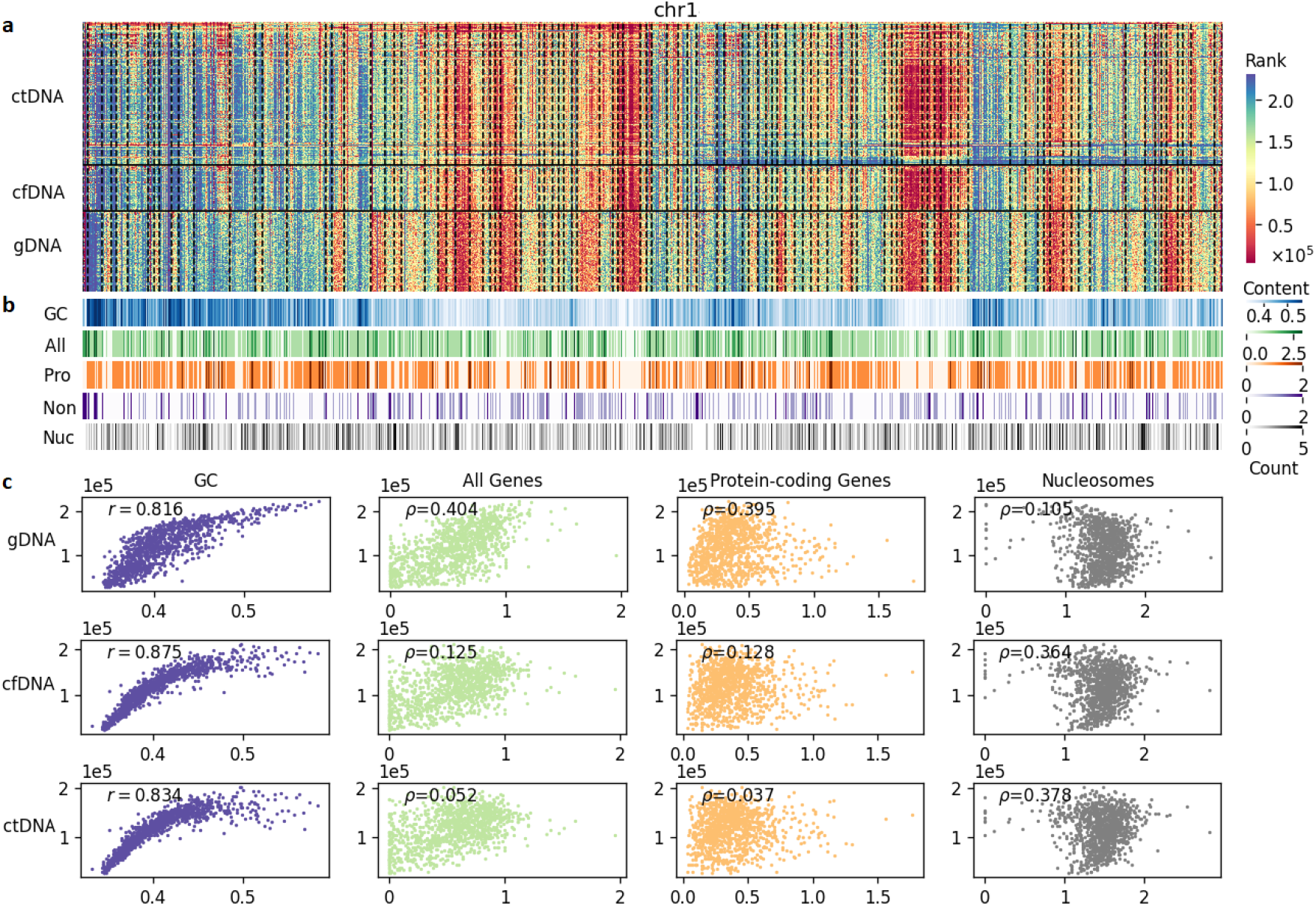
The coverage landscapes of gDNA, cfDNA and ctDNA. **(a)** Heatmap visualizing the rank-transformed coverage profiles of ctDNA, cfDNA and gDNA over 10kb windows at chr1. colour segments are evident, demonstrating the conserved patterns of coverage segmentation regardless of sample types. Dash lines indicate segment borders. Since there are too many gDNA (total=3,202) samples, here we only show 150 unselected gDNA samples to align with the numbers of other sample type. **(b)** Heatmap showing the GC-content (GC), gene density (All), protein-coding gene density (Pro), non-coding gene density (Non) and nucleosome density (Nuc) within each 10kb window at chr1. **(c)** Scatter plots of genome-wide coverage rank vs GC-content, (all) gene density, protein-coding gene density and nucleosome density in gDNA, cfDNA and ctDNA.

**Figure 3.**
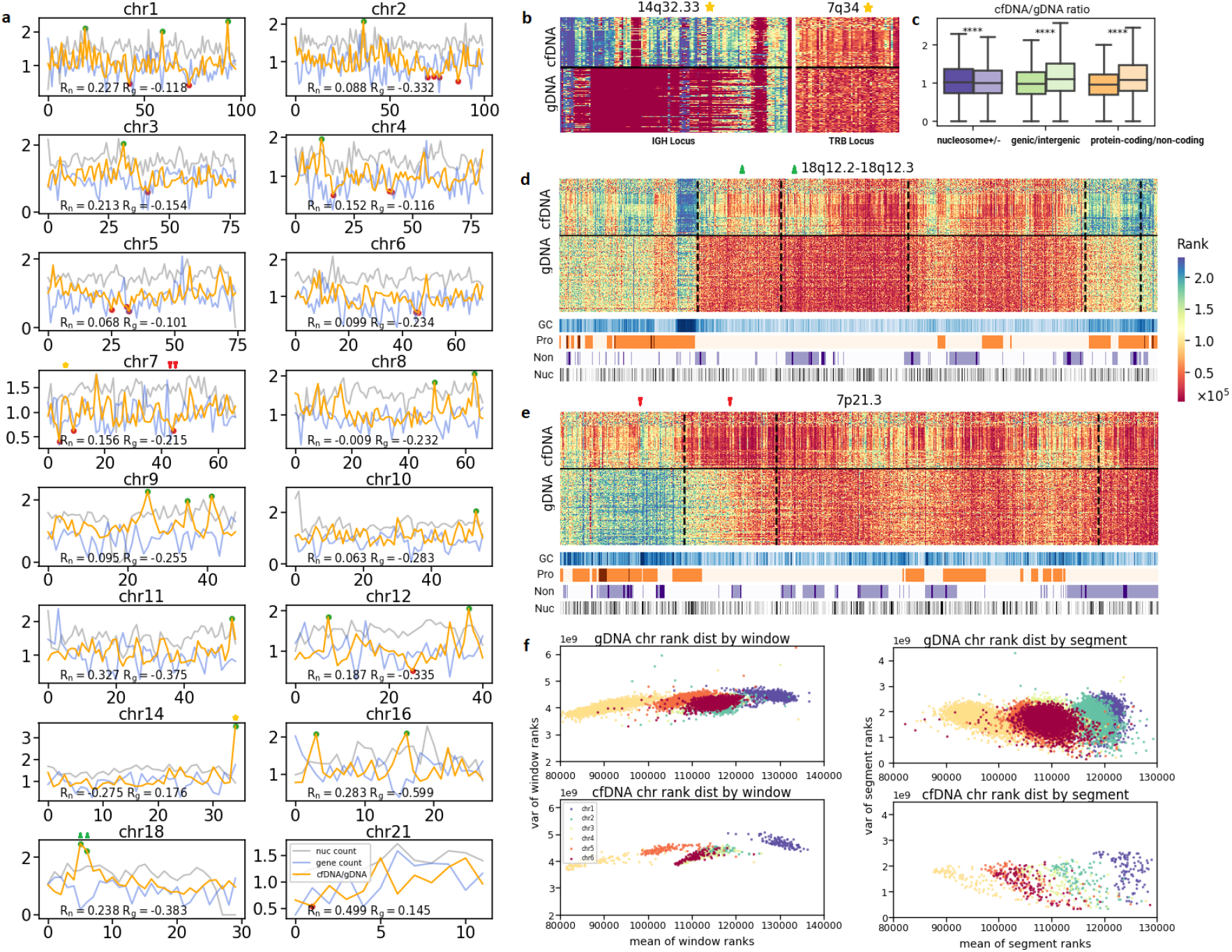
Differential coverage of cfDNA vs gDNA. **(a)** Line plots of segment coverage rank ratio of cfDNA vs gDNA (orange lines) in different chromosomes, plotted together with nucleosome density (nuc count, grey lines) and gene density (gene count, blue lines). The Pearson correlation coefficient between cfDNA/gDNA ratio and nucleosome or gene density (Rn, Rg) are shown for each chromosome. Top20 upregulated and top20 downregulated segments are highlighted in green and red circles, respectively. Chromosomes without top20 up/downregulated segments are not shown. A 3-fold coverage difference was observed in the last segment of chr14 (highlighted with star) mapping the IGH locus. The TRB locus at chr7 is also highlighted with star. Two typical cfDNA upregulated segments are highlited with green up-triangles, while two typical downregulated segments are highlighted with red down-triangles. **(b)** Heatmap visualization of the differential coverage at the IGH locus and the TRB locus. **(c)** The gDNA/cfDNA coverage ratio at windows mapping nucleosome positive or negative regions, genic or intergenic regions, and protein-coding genes or non-coding genes. Mann–Whitney–Wilcoxon tests were performed to determine whether the difference in ratio was significant. ****: p<10^−4^. **(d)** Heatmap showing typical examples of cfDNA upregulated segments (highlighted with green up-triangles), plotted together with GC-content (GC), protein-coding gene density (Pro), non-coding gene density (Non), nucleosome density (Nuc). Dash lines indicate segment borders. While gDNA is uniformly covered (evenly coloured with red), cfDNA is unevenly covered (blue, yellow, red intervals). **(e)** Heatmap showing typical examples of cfDNA downregulated segments (highlighted with red triangle), plotted together with GC-content (GC), protein-coding gene density (Pro), non-coding gene density (Non), nucleosome density (Nuc). Dash lines indicate segment borders. **(f)** Scatter plots showing the coverage mean and variance for chromosomes 1-6, calculated using the genome-wide 10kb-windows or segment representation.

### Calculations of gene and nucleosome density

Gene annotations were obtained from gencode gff3 version 46. Stable nucleosome positions from healthy individuals were obtained from the GSE81314 study (20). For each 10kb window, the count of all genes/protein-coding genes/non-coding genes/nucleosomes covered by the window was calculated as density.

### Computing the borders of coverage segmentation

We observed the phenomenon of coverage segmentation, that similar levels of coverage ranks were shared among adjacent windows to form segments. To locate the segments we need to find the borders of each segment. The python package ruptures (21) was used to find the borders of the coverage segments for each chromosome, using the median coverage rank of the 3,202 1kGP gDNA samples at each 10kb window. Specifically, we chose binary segmentation as the search method, with minimum segment size set to 100 windows (i.e., ≥ 1Mbp). To allow automatic determination of number of segments (breakpoints), penalty was set to log(n) · *σ*, where n is the total number of windows for the chromosome, *σ* is the standard deviation of the window ranks of the chromosome.

### Identification of up/downregulated segments

We compared the median segment ranks of all ctDNA samples (n=362) vs all cfDNA samples (n=113) to get top20 upregulated or downregulated segments and performed gene ontology enrichment analysis on genes underlying these segments using the R package clusterProfiler (22). Segments from chr19 were excluded due to significant inter-dataset variations demonstrated above (Supplementary Figure 3). While significant (FDR<0.05) enrichment was found in top20 downregulated segments, no significant enrichment was found in top20 upregulated segments (Supplementary Tables 1-2).

Then we examined each cancer dataset to identify ctDNA frequently up/downregulated segments. The coverage ratio of each segment was calculated between the median of each cancer dataset versus the median of the condition-matched controls (if existed) or the total control (n=113), except for the four prostate cancer samples from EGAD00001005343 due to too small sample size. Segments with at least 10% upregulation or downregulation were listed for each cancer dataset. The intersection of all lists was defined as frequently up/downregulated segments. Under this criteria, no intersection was found for downregulated segments, while three segments were found frequently upregulated (Supplementary Table 3).

### Differentially covered genes

For each gene, coverage is defined as the mean rank of the windows covering the gene. Similar to that of low count genes, we discard genes with the lowest 10% mean coverage in ctDNA and cfDNA. We also discarded highly variable genes in cfDNA whose variance exceeds the top 10% variance of all windows. Then for each gene the log2 fold-change (log2FC) between ctDNA and cfDNA was calculated. Mann–Whitney–Wilcoxon tests with Bonferroni corrections were performed on the log2FC values. Genes with FDR≤0.01 and absolute log2FC≥0.5 were identified as differentially covered genes.

### Rank histogram and outlier detection

After the rank transformation each sample has 236,239 10kb-windows whose coverage ranked from 1 to 236,239. An eight-bin histogram (each bin sized 30,000) was computed for each chromosome describing the distribution of ranks for the chromosome. In the four control datasets, the maximum median standard deviation of segment ranks was 14766.6 (EGAD00001006237, n=38), hence the choice of the 30,000 bin size is roughly two standard deviations.

The outlier detection method only requires sufficient number of control samples (n ≥20) for training. We used the Elliptic Envelope algorithm (23) implemented in scikit-learn (24) to learn the control histogram distribution of each chromosome, with the expected contamination rate set to 0.05, indicating a low level of (5%) outlier is expected in the training data. We filtered out chr19 (due to significant inter-dataset variations mentioned above) and chromosomes with less than 20 segments (chr21, 22), resulting in nineteen chromosomes (chr1-18, chr20) with a total of 1,040 segments. Each model is specifically trained for a chromosome. Each model for each chromosome gives a prediction score of whether the given histogram is an inlier/outlier (0 for outlier, 1 for inlier). The prediction score for each chromosome is weighted according to the number of segments in the chromosome. For example, chr1 has 98 segments, chr2 has 100 segments, then the weights for chr1 and chr2 are 98/1040=0.094, 100/1040=0.096, respectively. We defined *β* as the weighted sum of all prediction scores, which is the proportion of segments that support an inlier prediction. For each sample, the proportion of inlier segments and the proportion of outlier segments sum to 1, thus the proportion of outlier segments for the sample is 1 − *β*.

We defined the outlier proportion cutoff as *α*, samples with an outlier proportion 1 − *β > α* are deemed significantly different from the controls. The outlier proportion cutoff *α* is determined by the wanted specificity (default=0.9). We calculate the outlier proportion value for each control sample and put into a sorted list *A*. The cutoff *α* is the *n*^*th*^ value in the sorted list *A*.

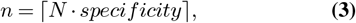

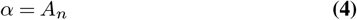

Where *N* is the number of control samples, and *n* is the ceil of the product of *N* and the wanted specificity. Test samples with outlier proportion exceeding the cutoff *α* are classified as cancerous.

### Benchmarks

For benchmarking, we chose ichorCNA (1) as an representative of copy-number-based methods, and LIQUORICE (9) as an representative of fragmentomics-based methods. LIQUORICE was developed based on samples from EGAD00001007080, one of the major dataset for our performance evaluation; ichorCNA is a popular tool for tumour content prediction and has been used in the EGAD00001007080 study.

For ichorCNA, we obtained the detailed ichorCNA results for EGAD00001007080 from the original study (9), and we used the default cutoff value 0.03 as cutoff for the maximum tumour content, i.e., classifying as cancerous if a sample with predicted maximum tumour content ≥0.03. Samples in EGAD00001006237 are early-stage lung adenocarcinoma (LUAD) samples, parameters recommended for low tumour content was used with the default 1Mb bins and 0.03 tumour content cutoff.

For LIQUORICE, Ewing-sarcoma-specific DHSs and Hematopoietic-specific DHSs were obtained from the original paper and used as region sets for EGAD00001007080. Since the Ewing-sarcoma-specific DHSs might not fit other types of sarcoma, to obtain optimized results, only Ewing-sarcoma and Ewing-sarcoma-like samples (n=206) and the matched controls (n=22) were used in the performance evaluation for LIQUORICE. LUAD-specific DHSs was obtained from ENCODE DNase Clusters specific to A549 LUAD cell line as instructed; LUAD-specific DHSs and Hematopoietic-specific DHSs were used as region sets for EGAD00001006237. Both datasets were then run with default parameters. For each sample, if the “Dip area: interpretation vs controls in same region set” or “Dip depth: interpretation vs controls in same region set” is flagged “Significantly stronger” or “Significantly weaker”, the sample was classified as cancerous. This setting allowed highest sensitivity and the best results in most cases, except causing zero specificity in the condition-unmatched task on EGAD00001007080. Therefore, to avoid zero specificity for the condition-unmatched task on EGAD00001007080 we classified samples flagging significantly different in both “Dip area: interpretation vs controls in same region set” AND “Dip depth: interpretation vs controls in same region set” as cancerous.

## Results

### Coverage landscape is conserved between gDNA and cfDNA

We first looked at the overall similarity of genome-wide coverage landscape between sample types. To integrate samples from different studies, we performed a rank transformation of the coverage profiles over 10kb windows. For each sample, windows were ranked based on their coverage, the higher the coverage the larger the rank (see Methods). The rank transformation showed advantages over raw coverage fraction, Z-score standardization and min-max normalization on dealing with batch effects (Supplementary Figure 1). Post-transform prinicple component analysis demonstrated substantial aggregation of control cfDNA samples from multiple datasets, and separation of ctDNA samples with (condition-matched) control cfDNA samples (Supplementary Figure 2). All chromosomes demonstrated inter-dataset consistency, except chr19 being highly variable within and between datasets (Supplementary Figure 3), corresponding to previous findings that intra-species variation of human common single-nucleotide polymorphisms was most prevalent on chr19 (25).

The rank-transformed coverage profiles were visualized with heatmaps and compared between gDNA, cfDNA and ctDNA (Figure 2a). From the heatmaps we observed the aggregation of windows into segments (i.e., adjacent windows share similar colours), indicating neighboring windows tend to present similar coverage (ranks). The patterns of segmentation seemed to be largely conserved even between gDNA and cfDNA, allowing the use of segments to represent the genome-wide windows for noise and dimensionality reduction. We computed the borders of these segments throughout the genome using the 3,202 gDNA samples from 1kGP (see Methods), resulting in 1,090 segments of variable length (≥1Mb) across the genome (Figure 2a, dashed vertical lines).

We identified major contributing factors to coverage (Figure 2b-c). Note that to preserve the original coverage landscape and to investigate the contributions of multiple factors that could be confounded by GC-content, GC-correction was not applied. Consistent with previous findings (26), GC-content showed a strong positive correlation with the 10kb-window coverage (R=0.816). We also confirmed the confounding effect of GC-content (Supplementary Figure 4) as it showed positive correlation with gene density (R=0.537) and weak negative correlation with nucleosome density (R=-0.197). Interestingly, significant partial correlation was found between gene density and gDNA coverage (*ρ*=0.404, GC-controlled) but not between gene density and cfDNA coverage (*ρ*=0.13, GC-controlled). The contribution of gene density to gDNA coverage was mainly by protein-coding genes (*ρ*=0.395, GC-controlled) - segments enriched with protein-coding genes seemed to present higher coverage, while segments with sparse protein-coding genes are less covered (Figure 2a-b). However in cfDNA, nuclosome density (*ρ*=0.364, GC-controlled) instead was the contributor of coverage second to GC-content. Our results indicated that apart from GC-content, the positioning of protein-coding genes affects gDNA coverage, whereas nucleosome positioning affects cfDNA coverage.

**Figure 4.**
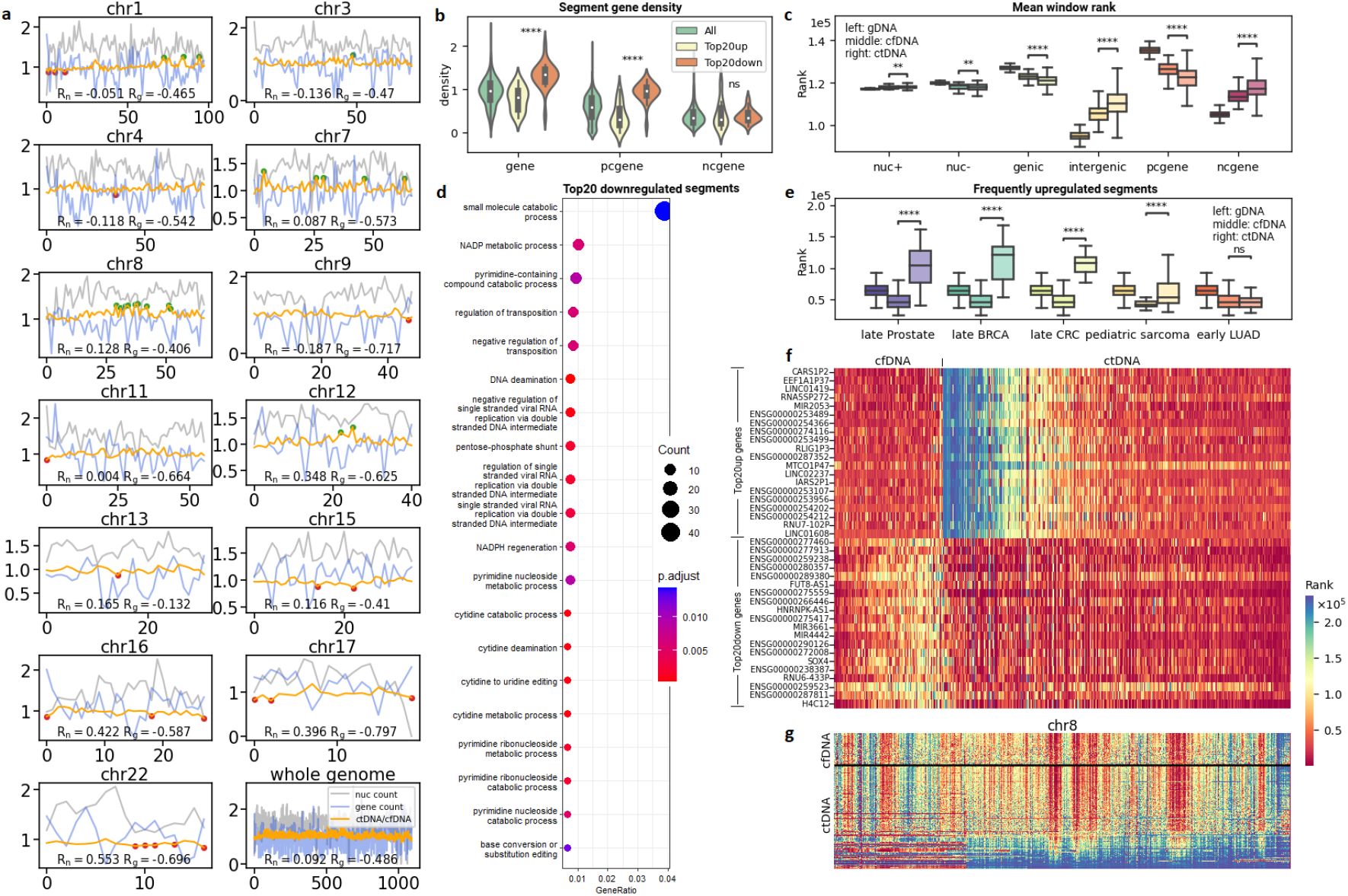
Differential coverage of ctDNA vs cfDNA. **(a)** Line plots of coverage rank ratio of ctDNA vs cfDNA (orange lines) on segments within different chromosomes, plotted together with nucleosome density (nuc count, grey lines) and gene density (gene count, blue lines). The Pearson correlation coefficient between ctDNA/cfDNA ratio and nucleosome or gene density (Rn, Rg) are shown for each chromosome. Top20 upregulated and top20 downregulated segments are highlighted in green and red circles, respectively. Chromosomes without top20 up/downregulated segments are not shown. Upregulated segments are significantly enriched at chr8. **(b)** Violin plot of gene density, protein-coding gene (pcgene) density and non-coding gene (ncgene) density in all segments, top20 upregulated segments and top20 downregulated segments. **(c)** A comparison of gDNA (n=3,202), cfDNA (n=113) and ctDNA (n=362) mean coverage at windows mapping nucleosome positive (short-named as nuc+) regions, nucleosome negative (nuc-) regions, genic regions, intergenic regions, protein-coding genes (pcgene) or non-coding genes (ncgene). Mann–Whitney–Wilcoxon tests were performed between ctDNA and cfDNA to determine whether differential coverage was significant. **: p<10^−2^, ****: p<10^−4^. **(d)** Significantly enriched gene ontology terms of biological process from genes within the top20 downregulated segments, FDR<0.05. **(e)** A comparison of gDNA, cfDNA and ctDNA mean coverage at three ctDNA frequently upregulated segments on multiple cancer datasets. **(f)** Differentially covered genes (FDR*≤*0.01 and log2FC*≥*0.5), with top20 upregulated and top20 downregulated shown. **(g)** Heatmap visualization on coverage landscape of ctDNA and cfDNA at chr8. The distinctive convergence of segment coverage is evident in ctDNA samples at the bottom.

### cfDNA vs gDNA differential coverage at immune-receptor loci, intergenic regions and non-coding genes

Next, we compared the segment coverage profile of cfDNA (n=113) and gDNA (n=3,202). For each sample, the coverage of each segment is defined as the mean of window ranks within the segment. We identified top20 cfDNA (versus gDNA) upregulated or downregulated regions, respectively (Figure 3a). We observed a 3-fold upregulation of cfDNA at the segment mapping the IGH locus located 14q32.33 (Figure 3a-b; Supplementary Figure 5). We then examined whether differential coverage also occur in the TRB locus located at 7q34. Although not as strong as that of the IGH locus, a 32% increase was found in cfDNA on the segment mapping the TRB locus (Figure 3a-b). The IGH locus is rearranged in B cells and partially rearranged in T cells (27), meaning a large part of the locus is lost in lymphocytes, which account for about 80% of total PBMCs. This explains the extremely low coverage at the IGH locus in the gDNA samples, because the cell sources of 1kGP gDNA samples are PBMCs. Reversely, from the dramatic IGH upregulation in cfDNA we can infer that lymphocytes are not the major cell sources of cfDNA. Indeed, neutrophils are the most abundant leukocytes, and their short lifespan will even add to their contribution to dead cells in the blood- the source of cfDNA. Compared to the broad IGH partial rearrangement in T cells, the TRB rearrangement in B cells from healthy individuals is less prevalent and restricted to T-cell receptor (TCR) *δ* genes (28). However, a subset of circulating neutrophils (5-8%) is known to express TCRs (29) and thus having rearranged TRB locus. As a result, greater coverage difference was observed in the IGH locus, compared to the TRB locus.

**Figure 5.**
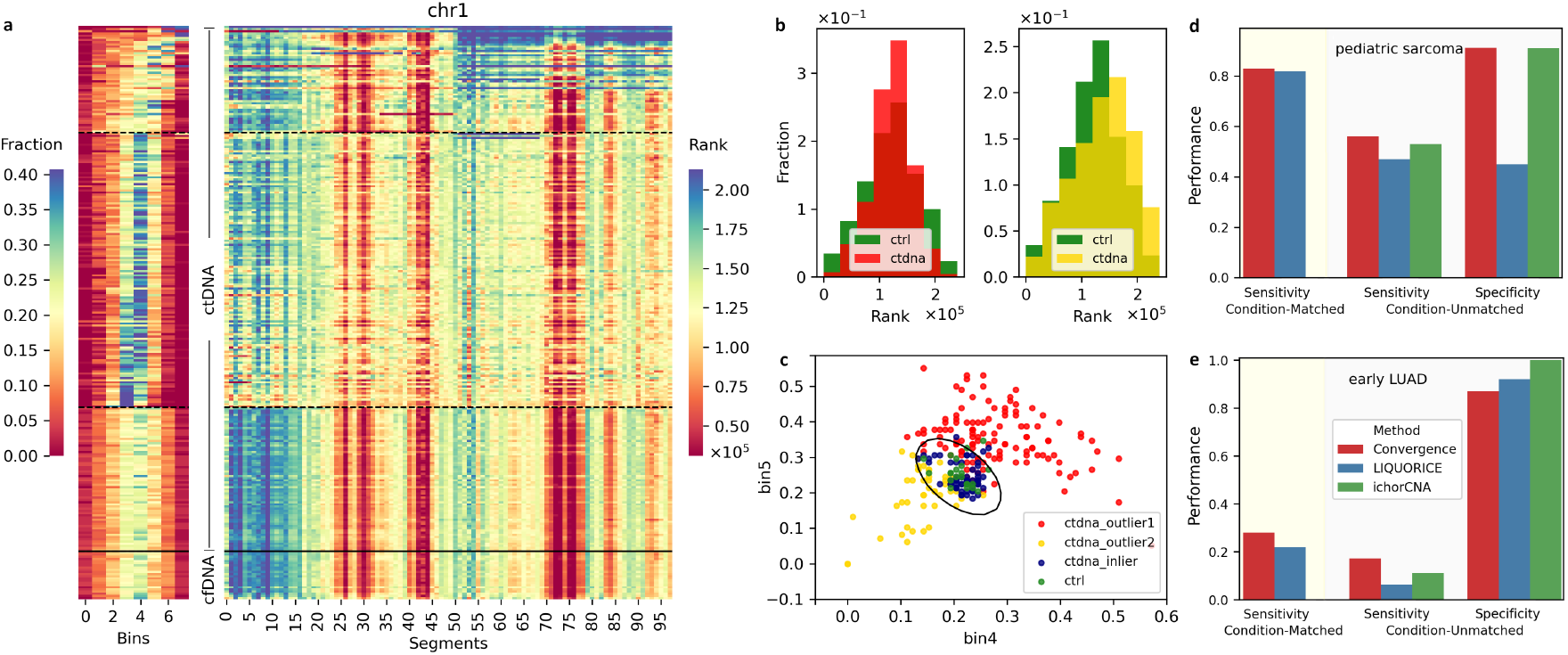
Outlier detection on coverage profiles, exampled with chr1. **(a)** Heatmap visualization of an 8-bin histogram (left panel) of segment coverage rank and the original segment coverage rank (right panel) at chr1 of ctDNA from pediatric sarcoma samples (n=241) versus condition-matched control cfDNA (n=22). Solid lines separate control and cancer samples. Dashed lines separate cancer samples exhibiting different directions of coverage convergence into three parts: the top part shows convergence from middle bins towoards edge bins and corresponds to the right panel of Figure 5b; the middle part shows convergence from edge bins towards middle bins and correspondes to the left panel of Figure 5b; the bottom part corresponds to no appearint convergence. **(b)** Differential rank histogram of sarcoma-derived ctDNA vs condition-matched control cfDNA. Convergence of segment coverage was observed, either towards the middle-bins (left panel) or towards the end-bins (right panel). **(c)** Scatter plot of two middle bins of the 8-bin histogram (bin4 and bin5) of the ctDNA outliers (highlighted in red and yellow colours, corresponding with Figure 5b) detected by Elliptic Envelope. Black oval indicates the decision boundary of the Elliptic Envelope model. **(d)** Performances of the proposed convergence-based cancer detection method with benchmarks on pediatric sarcoma (n=241) and matched-control (n=22), detailed in Table 1. **(e)** Performances of the proposed convergence-based cancer detection method with benchmarks on early lung adenocarcinoma (early-LUAD, n=36) and matched-control (n=38), detailed in Table 1.

Besides immune-receptor loci, cfDNA was found significantly upregulated in intergenic regions and non-coding genes (Figure 3c), compared to genic regions and protein-coding genes, respectively. The cfDNA/gDNA coverage ratio was not affected by GC-content (R=-0.067; Supplementary Figure 6). Gene density negatively correlated with cfDNA/gDNA ratio in most chromosomes (Figure 3a; Supplementary Figure 7). Upregulation of cfDNA was typically observed in segments without protein-coding genes but non-coding genes; gDNA seemed to be uniformly covered in these segments, while apparent ups and downs (reflected by colour changes in heatmap) present in the coverage of cfDNA (Figure 3d, two cfDNA upregulated segments highlighted with green triangles). Meanwhile, cfDNA downregulation was typically observed in protein-coding regions (Figure 3e, two cfDNA downregulated segments highlighted with red triangles). The differential coverage between cfDNA and gDNA reflects the distinct activities (e.g. rearrangement, apoptosis) happen within different cell sources.

Despite significant differential coverage in local regions, the distribution of coverage is largely conserved between cfDNA and gDNA at chromosome level (Figure 3f). The mean and variance of each chromosome distinguish from one another; meanwhile, the mean and variance of each chromosome remain relatively constant between cfDNA and gDNA. While levels of variances differ significantly between chromosomes with the genome-wide windows, the 1,090 segments well adjusts them to similar levels.

### Distinctive convergence of segment coverage in ctDNA

We then compared segment coverage of ctDNA vs cfDNA to identify potential tumour-induced changes. Similarly, we identified top20 ctDNA (versus cfDNA) upregulated or downregulated segments (Figure 4a; Supplementary Tables 1-2). Significant differences in protein-coding gene density were found between the top20 upregulated and the top20 downregulated segments - the downregulated segments were enriched with protein-coding genes while the upregulated ones were sparse (Figure 4b). Moreover, we observed a further drop of coverages at genic regions or protein-coding genes and a further rise in coverages at intergenic regions or non-coding genes (Figure 4c), compared to the cfDNA vs gDNA differential coverage shown above.

This suggested a decreased contribution by normal cells due to the replacement by other cell sources with distinct activities, such as the tumour. We also observed a stronger negative correlation (R=-0.486, whole genome; Figure 4a; Supplementary Figure 8) between ctDNA/cfDNA ratio and gene density, compared to the negative correlation between cfDNA/gDNA ratio and gene density (R=-0.233, whole genome; Supplementary Figure 7). The top20 downregulated segments were significantly (FDR<0.05) enriched with genes associated with biological processes involving DNA/RNA mutation, nucleotide metabolism and NADPH metabolism (Figure 4d), which are closely related to cancers (30–32). Due to low gene density, no significant enrichment was found in the top20 upregulated segments. However, we identified three frequently upregulated segments with at least 10% increase in median coverage in all cancer datasets compared to (condition-matched) controls (see Methods; Figure 4e; Supplementary Table 3). The frequently upregulated segments significantly (p<10^−4^) distinguished ctDNA (prostate, breast, colorectal cancers and pediatric sarcoma) from cfDNA, except for early-stage lung cancer, which showed a non-significant trend (p=0.146). Similarly to the concept of differentially expressed genes, here we defined differentially covered genes (see Methods). We identified 797 differentially covered genes with FDR≤0.01 and log2FC≥0.5 (Figure 4f; Supplementary Table 4), in which 549 were upregulated and 248 were downregulated. Enrichment analysis on the downregulated genes identified significant associations with RNA splicing, epithelial proliferation and protein-kinase activities (Supplementary Figure 9). Similar to the top upregulated segments, no significant enrichment was found in the upregulated genes.

Of note, we observed a distinctive feature of ctDNA – the convergence of segment coverage during progression of cancer. A typical example is seen in chr8, where ctDNA upregulated segments accumulated in the long arm (Figure 4a, 4g), consistent with the fact that chr8 gain is common in cancers (33, 34). The degree of segment convergence appeared to be positively associated with cancer stages, with a gradual loss of correlation to normal (control) segment values as cancer progress (Supplementary Figure 10). While early-stage cancers may not show visible convergence on heatmap, late-stage cancers typically converge into a small number of segments, which are visually compelling (Supplementary Figure 11).

### Measuring segment convergence for the detection of cancer

Finally, we examined the possibility of coverage-based cancer detection by calculating the segment convergence in chromosomes. From a chromosomal point of view, convergences can happen in any segments towards either directions – converging to higher or lower coverage ranks. Since the ranks are calculated genome-wide, changes of segment rank distribution in one chromosome would affect the segments in other chromosomes. The segment rank distribution can be represented as a histogram, which partitions the ranks into bins. Figure 5 shows the comparison of an 8-bin segment rank histogram (alongside the segment heatmap) between ctDNA (pediatric sarcoma, n=241) and condition-matched control cfDNA (n=22) on chr1. From the comparative histogram, we observed a diverse spectrum of changes that could happen in ctDNA. In the most extreme cases, the whole chromosome or an arm converged to the highest or lowest bin; in most cases, segments belonging to the edge bins converged to the middle bins (Figures 5a-b).

Considering the position and direction of convergence is random, instead of looking for characteristics of the cancer samples, we transformed the question into a task of outlier detection. We assumed that the segment rank histograms of control cfDNA samples follow a multivariate Gaussian distribution. For each chromosome, we computed the 8-bin histogram, resulting in an 8-dim vector for each sample. The shape of an multi-dimensional Gaussian distribution is an ellipsoid, and we used the Elliptic Envelope (23) algorithm to first learn a model of the control elliptic distribution. Any points outside the ellipsoid were deemed outliers (Figure 5c). The model then can be used to predict whether a test vector is outlier. For each test sample, we weighted and summarized the prediction results for all chromosomes (see Methods). When the proportion of segments flagged as outliers exceeds a cutoff (see Methods), the test sample is deemed significantly different from controls.

The performances were assessed for two scenarios: prediction with or without condition-matched controls. Condition-matched controls are known control cfDNA samples sequenced together or under the same condition with the ctDNA samples. When condition-matched controls were not available, condition-unmatched controls (i.e., controls from other studies) were used for training; sensitivity and specificity were assessed. Table 1 shows details of the performances for both scenarios (highlighted in red), with benchmarks from state-of-the-art fragmentomic-based (9) or copy-number-based (1) methods. For the condition-matched tasks (Figures 5d-e; Table 1), the convergence-based method achieved 83% sensitivity on pediatric sarcomas (n=241) and 28% sensitivity on early lung cancers (n=36), outperforming benchmark. For the non-ideal, condition-unmatched tasks (Figures 5d-e; Table 1), the convergence-based method achieved 56% sensitivity on pediatric sarcoma (n=241), 17% sensitivity on early lung cancers (n=36), 95-100% sensitivity on late-stage cancers (n=85, from four datasets) with an overall 91% specificity on all controls (n=113, from four datasets). Note that the control distribution still vary between studies, likely due to a variety of factors such as sequencing platform, experiment protocols and sequencing depth, although not as strong as the differences between control and most cancer cases (Supplementary Figures 12-15). Therefore, condition-unmatched predictions should be interpreted with caution.

## Discussion

In this study, we systemically compared genome-wide coverage profiles between gDNA samples from 1kGP (n=3,202) and cfDNA samples of healthy individuals (n=113) and cancer patients (n=362). We observed the phenomenon of coverage segmentation which was largely conserved among gDNA and cfDNA, where adjacent genomic regions share similar levels of coverage, forming segments that correspond to GC-content and protein-coding gene density or nucleosome density. We identified differential cfDNA vs gDNA coverage at immune-receptor locus, intergenic regions and non-coding genes, which were associated with differential genome activities from different cell sources. We also discovered an intensified coverage drop in genic regions or protein-coding genes and rise in intergenic regions or non-coding genes in ctDNA, indicating the increased proportion of tumour-derived DNA in the plasma. Moreover, we found the convergence of segment coverage as a distinctive characteristic during cancer development and progression. We demonstrated as a proof-of-concept an outlier-detection method to measure the segment convergence for cancer screening without using any cancer data for training, i.e., only trains on control cfDNA data. Our method achieved versatile performances and good generalizability to multiple independent datasets on both condition-matched and condition-unmatched tasks.

We uncovered biological insights hidden beneath coverage. Despite well known effects of GC-content to sequencing coverage, differential coverage caused by differences in protein-coding/non-coding genes or genic/intergenic regions were rarely discussed. Like expression profiles, our results demonstrated that the genome-wide coverage profiles are cell-type specific. Lymphocyte is the major cell type of PBMC, whereas neutrophil is the most abundant leukocyte in blood. Non-lymphocyte cells have intact IGH locus while lymphocytes have partial or full rearrangements. As a result, coverage at the IGH locus has a 3-fold difference between PBMC-derived gDNA and plasma cfDNA. Since the lymphocyte count is strongly associated with health status, the IGH coverage which is proportional to lymphocyte-derived cfDNA, could be further utilized as an indicator of diseases.

Several processing steps are key to the coverage profiling analyses: 1) the rank transformation of the coverage fraction, which allows integration of data from different studies. Although not commonly seen in DNA analysis, the rank transformation has been used in handling single-cell expression counts as a non-parametric method that circumvents the need of normalization (35). 2) The computation of coverage segments to represent the genome-wide coverage windows, which extracts important features and de-noises the data. 3) Histogram binning of chromosomal segment coverage, which allows further feature selection. 4) Outlier detection, to detect outliers based on a known control distribution. This removes the dependence on cancer samples for training, which is considered more generalizable than conventional binary classifiers that requires cancer samples for training, because cancers are too heterogeneous that could not be fully explained by even thousands of samples.

Finally, to demonstrate the feasibility of using extracted features from genome-wide coverage profiles to detect early cancers, we chose the Elliptic Envelope algorithm (23) to perform outlier detection. Other outlier/novelty detection approaches or algorithms could also fit the task, and may further improve the performances.

## Data and Code Availability

All data used within this study can be retrieved from public databases with accession number listed in Table 1. Source code for this study is freely available at https://github.com/deepomicslab/Genome-wide-coverage-profiling.

## Acknowledgments

Not applicable.

