## Supplementary Figures 1-15 for "Coverage landscape of the human genome in nucleus DNA and cell-free DNA"

**Note: as mentioned in the main text, for the ease of expression, we use **ctDNA** to represent the entire cfDNA sample drawn from cancer patients, **cfDNA** to represent the entire cfDNA sample from healthy controls.

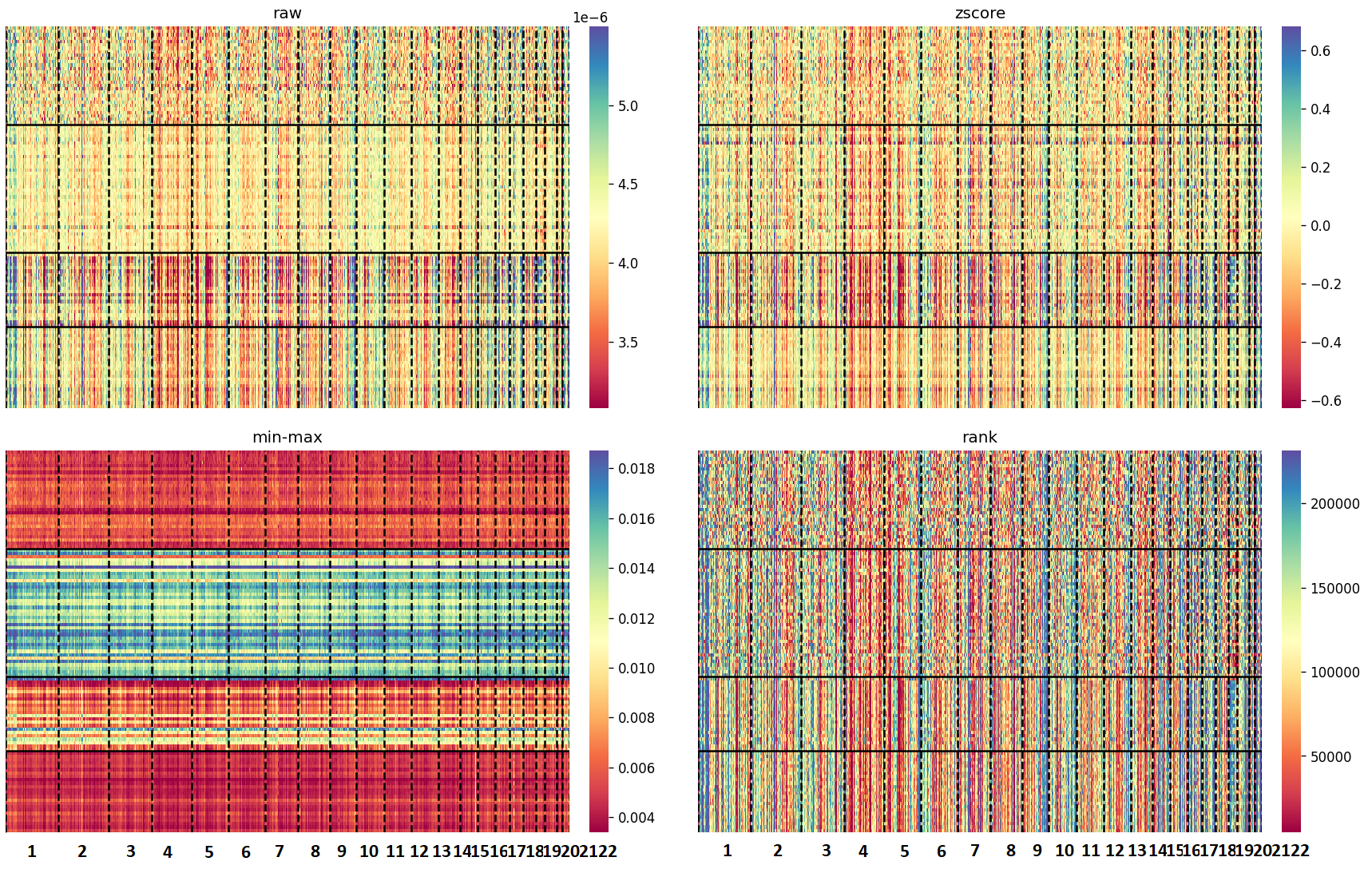

Supplementary Figure 1. Genome-wide 10kb windows (columns) of raw coverage fraction, z-score standardization, min-max normalization and rank transformation on four control cfDNA datasets (rows). Solid horizontal lines separate the datasets; dashed vertical lines separate chromosomes.

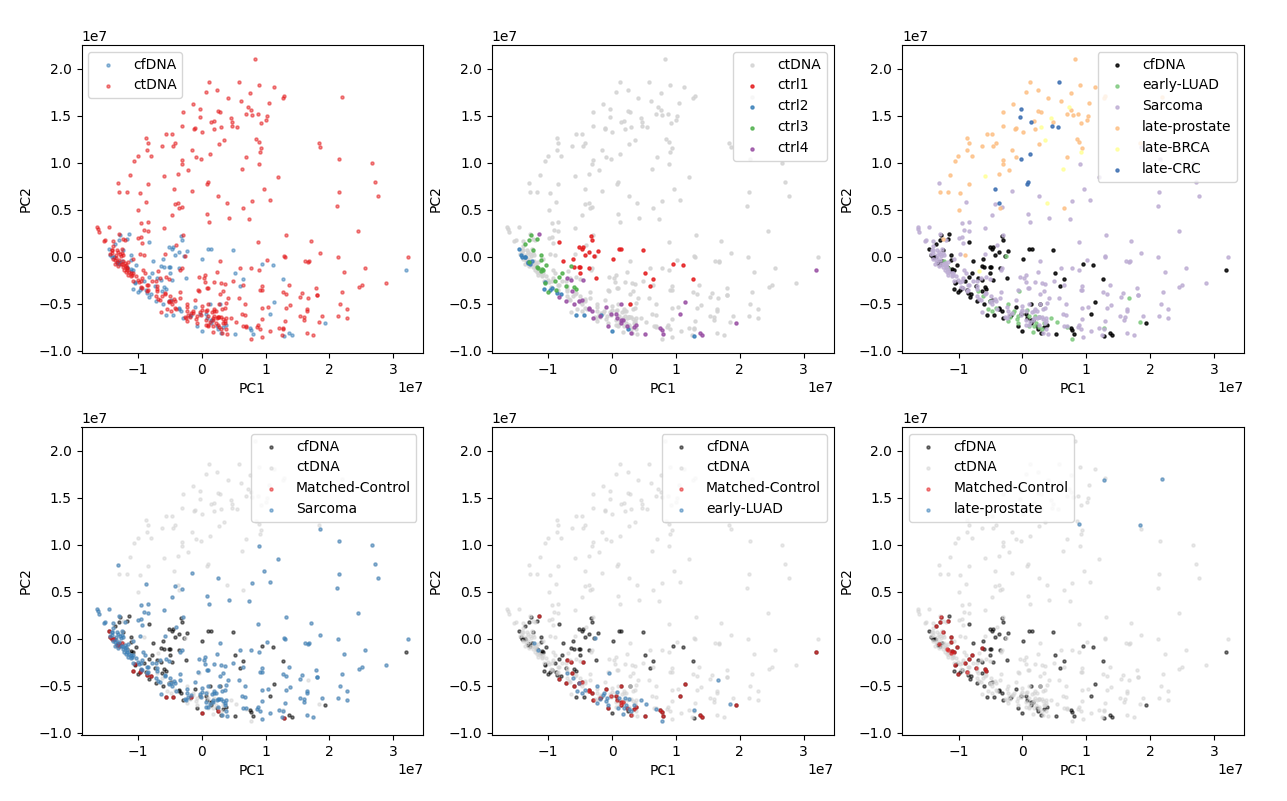

Supplementary Figure 2. Principle component analysis plots of post-transform data. Top left: all cfDNA (n=113) versus all ctDNA (n=362). Top middle: positions of samples from four control cfDNA datasets (ctrl1-4). Top right: positions of samples from multiple ctDNA datasets, note that the entry for late-prostate is a combination of two datasets EGAD00001008460 (n=60) and EGAD00001005343 (n=4). Bottom left: positions of ctDNA and cfDNA samples from EGAD00001007080, including pediatric sarcoma (n=241) and condition-matched controls (n=22). Bottom middle: positions of ctDNA and cfDNA samples from EGAD00001006237, including early lung adenocarcinoma (early-LUAD, n=36) and condition-matched controls (n=38). Bottom right: positions of ctDNA and cfDNA samples from EGAD00001005343, including late-prostate cancers (n=4) and condition-matched controls (n=24).

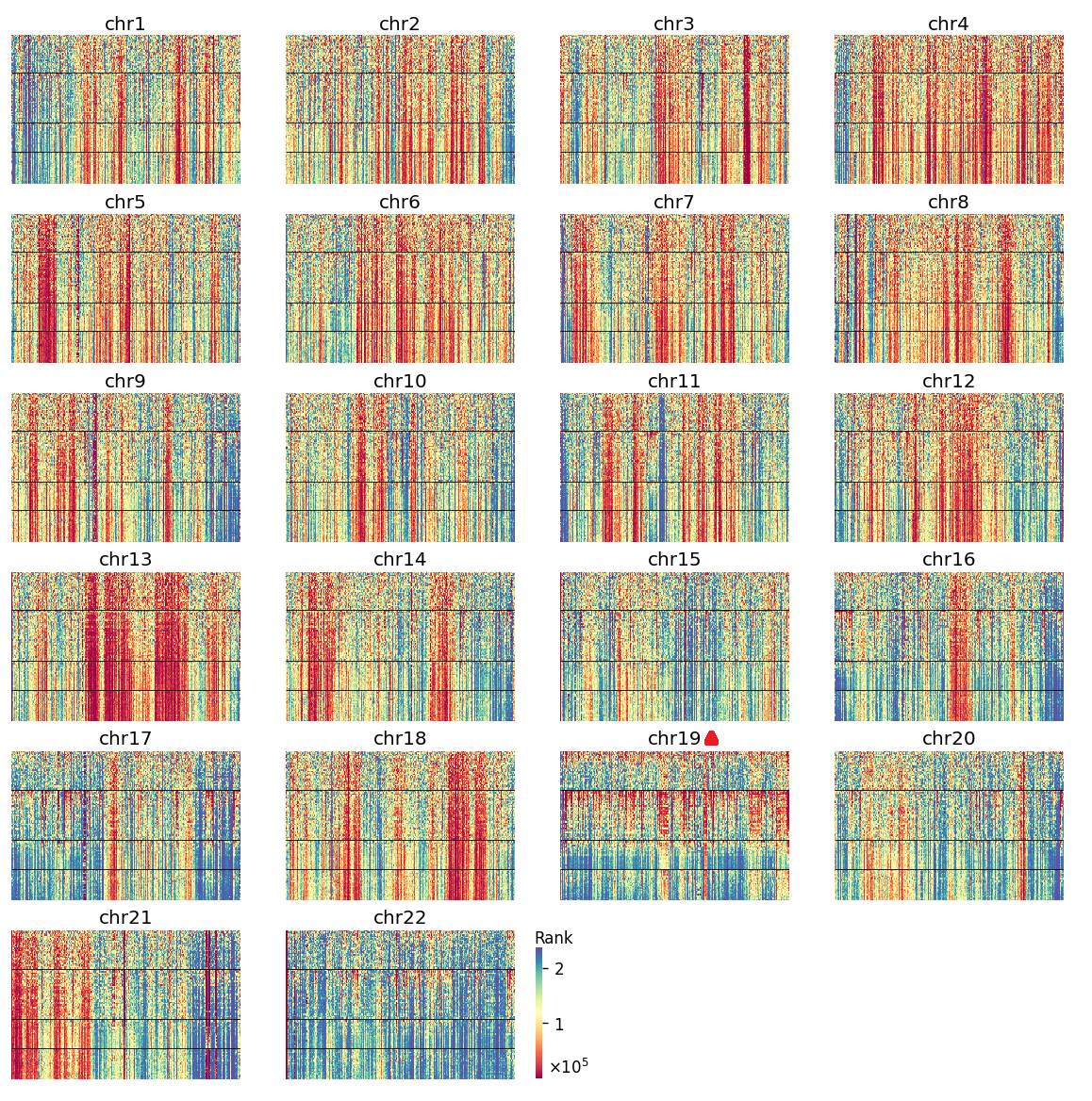

Supplementary Figure 3. Chromosomal view of coverage ranks over 10kb windows (columns) on four control cfDNA datasets (rows), separated by horizontal lines. Overall consistency was observed, except in chr19, where inter-dataset and intra-dataset variation was evident.

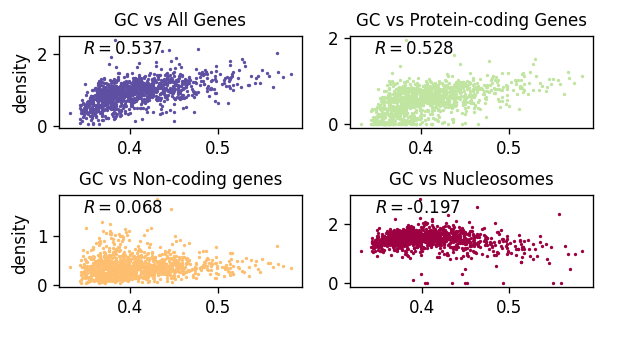

Supplementary Figure 4. Confounding effects of GC-content. Pearson correlation coefficients are shown between GC vs gene density (all genes), protein-coding gene density, non-coding gene density and nucleosome density.

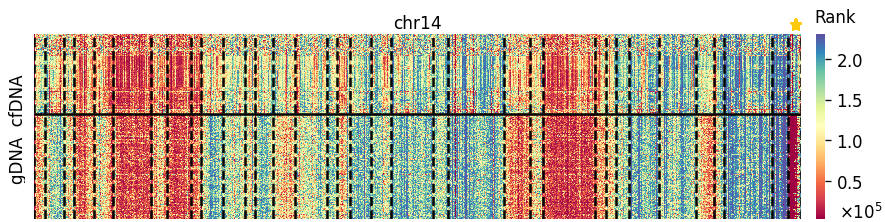

Supplementary Figure 5. Coverage heatmap of cfDNA vs gDNA on chromosome 14. A 3-fold difference found in the end segment, mapping the IGH locus located at 14q32.33, highlighted with star as related to main Figure 3a-b. Solid horizontal line separates cfDNA and gDNA samples; dashed lines indicate segment borders.

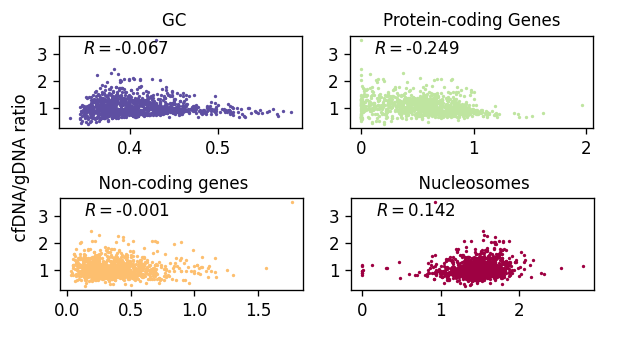

Supplementary Figure 6. Pearson correlation analysis between cfDNA/gDNA ratio and GC-content, protein-coding gene density, non-coding gene density and nucleosome density.

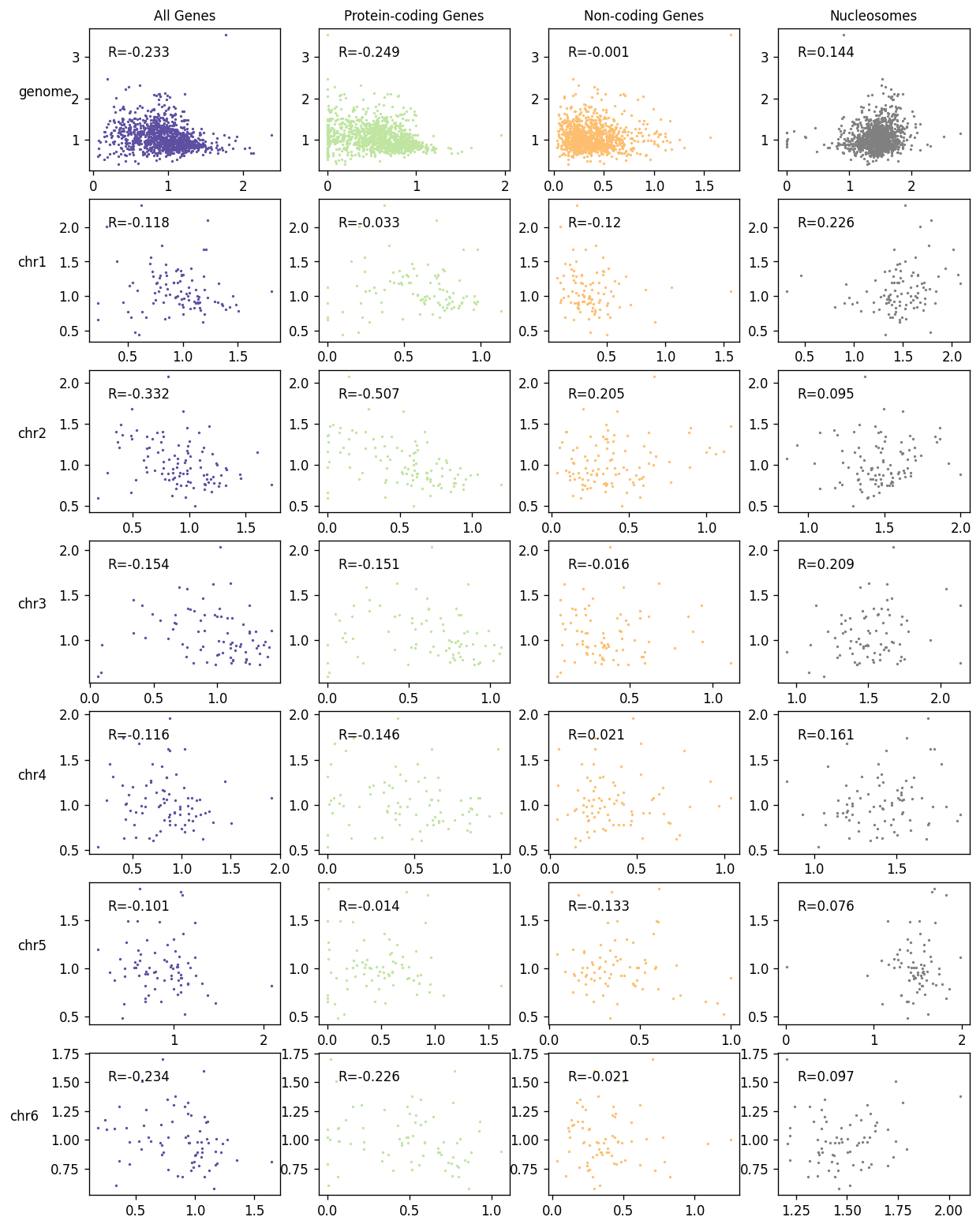

Supplementary Figure 7. Pearson correlation analysis between cfDNA/gDNA ratio and gene density, protein-coding gene density, non-coding gene density and nucleosome density at whole-genome level and chromosomes 1-6.

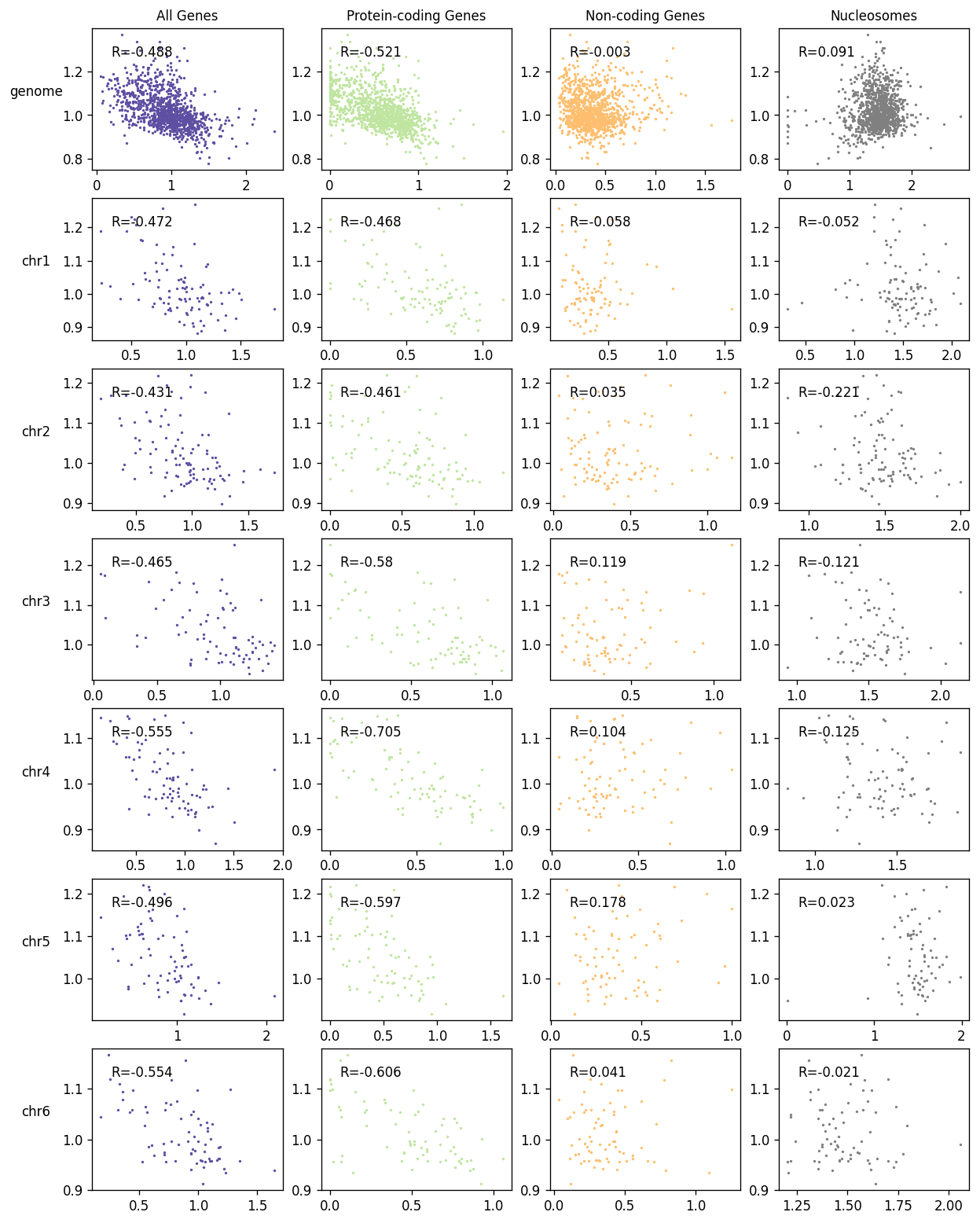

Supplementary Figure 8. Pearson correlation analysis between ctDNA/cfDNA ratio and gene density, protein-coding gene density, non-coding gene density and nucleosome density at whole-genome level and chromosomes 1-6.

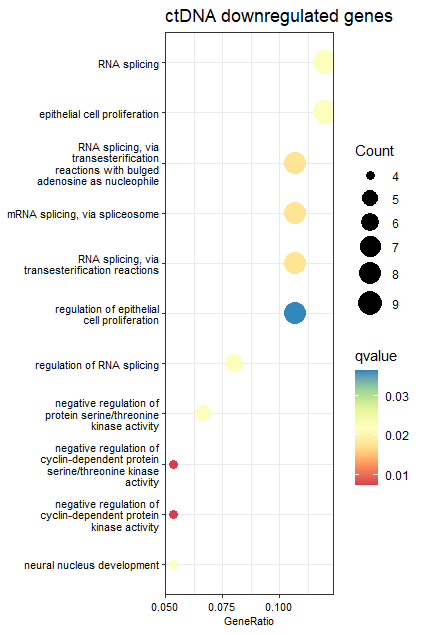

Supplementary Figure 9. Enrichment analysis (FDR<=0.05) of ctDNA (versus control cfDNA) downregulated genes. No significant enrichment was found for ctDNA upregulated genes.

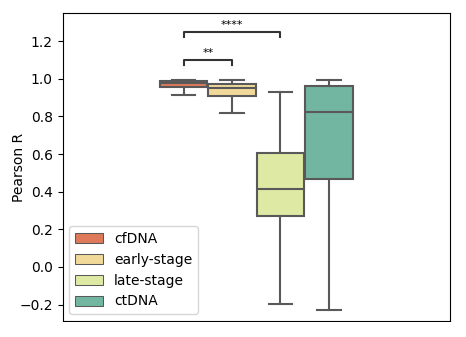

Supplementary Figure 10. Boxplots of correlation coefficients for the 1,090 segments between the median of 113 control cfDNA samples and each sample of cfDNA (control, n=113), early-stage (n=36), late-stage (n=85) and all ctDNA (n=362).

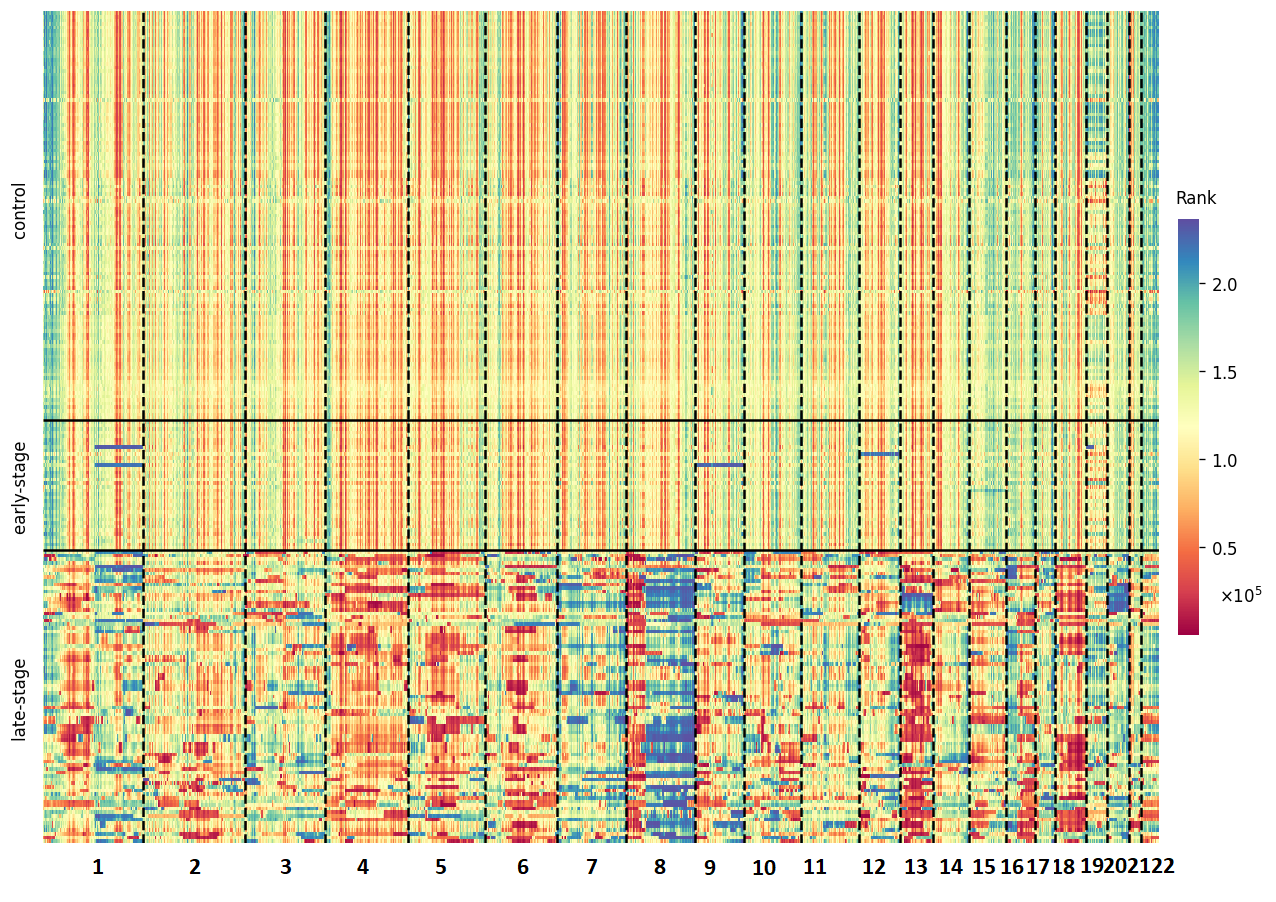

Supplementary Figure 11. A comparison of segment convergence in cfDNA (control, n=113), early-stage (n=36) and late-stage (n=85) ctDNA.

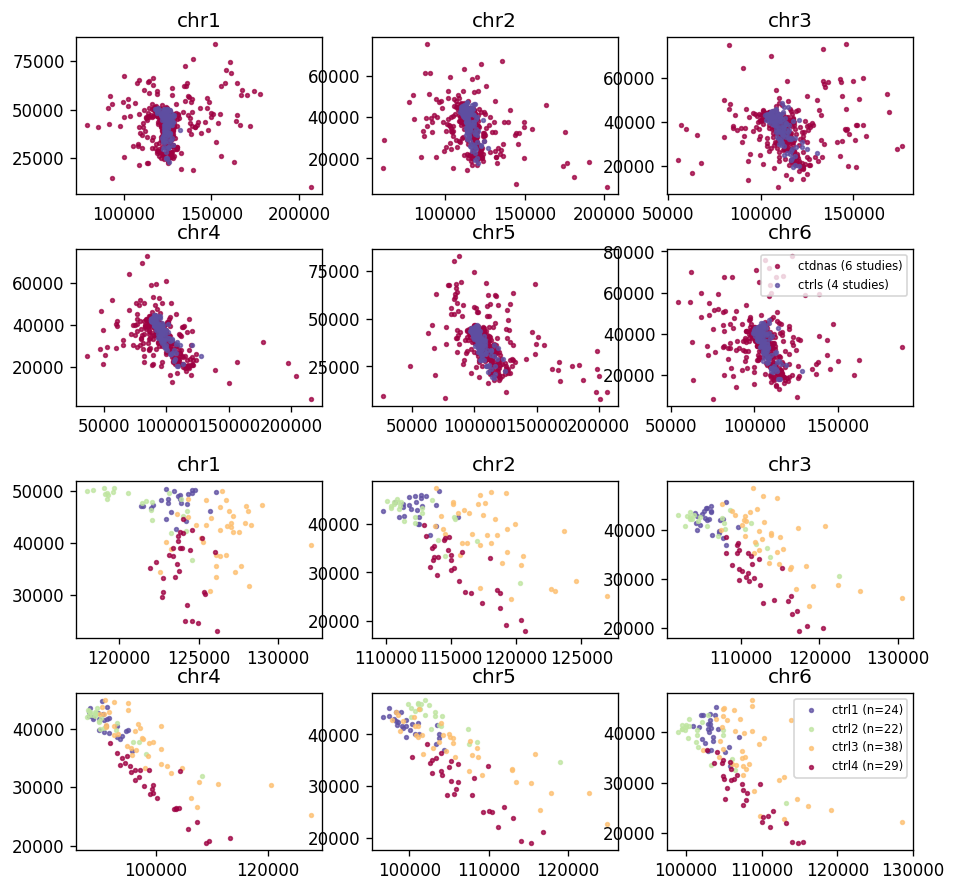

Supplementary Figure 12. Mean (x-axis) and standard deviation (y-axis) of segment coverage rank in ctDNA (from six datasets, n=362) vs control cfDNA (from four control datasets), for chromosomes 1-6.

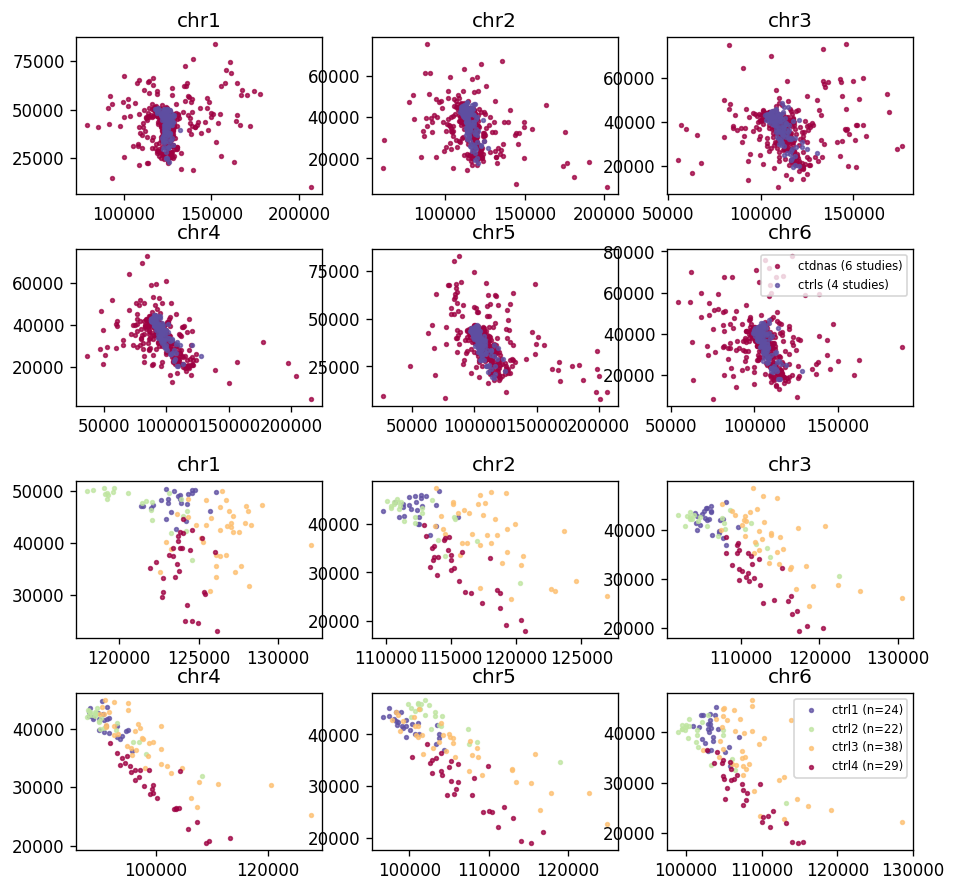

Supplementary Figure 13. Mean (x-axis) and standard deviation (y-axis) of segment coverage rank in the four control cfDNA datasets (ctrls1-4), for chromosomes 1-6.

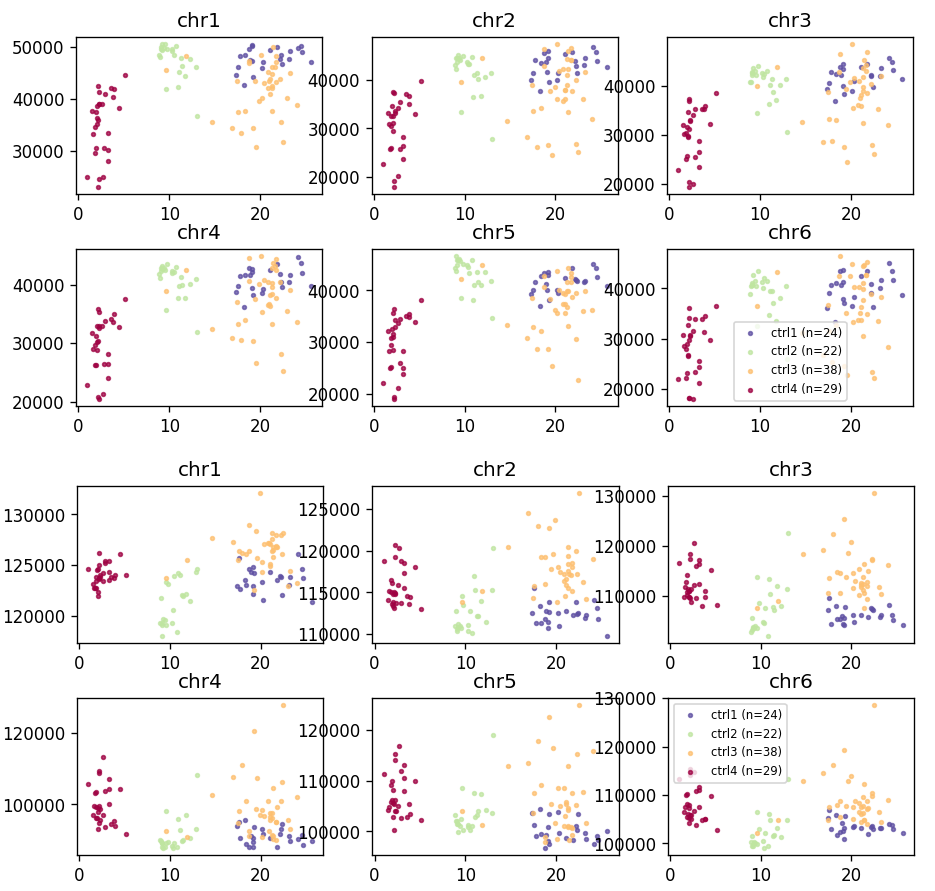

Supplementary Figure 14. Plots of mean sequencing depth (x-axis) versus mean segment coverage (y-axis) in the four control cfDNA datasets (ctrls1-4), for chromosomes 1-6.

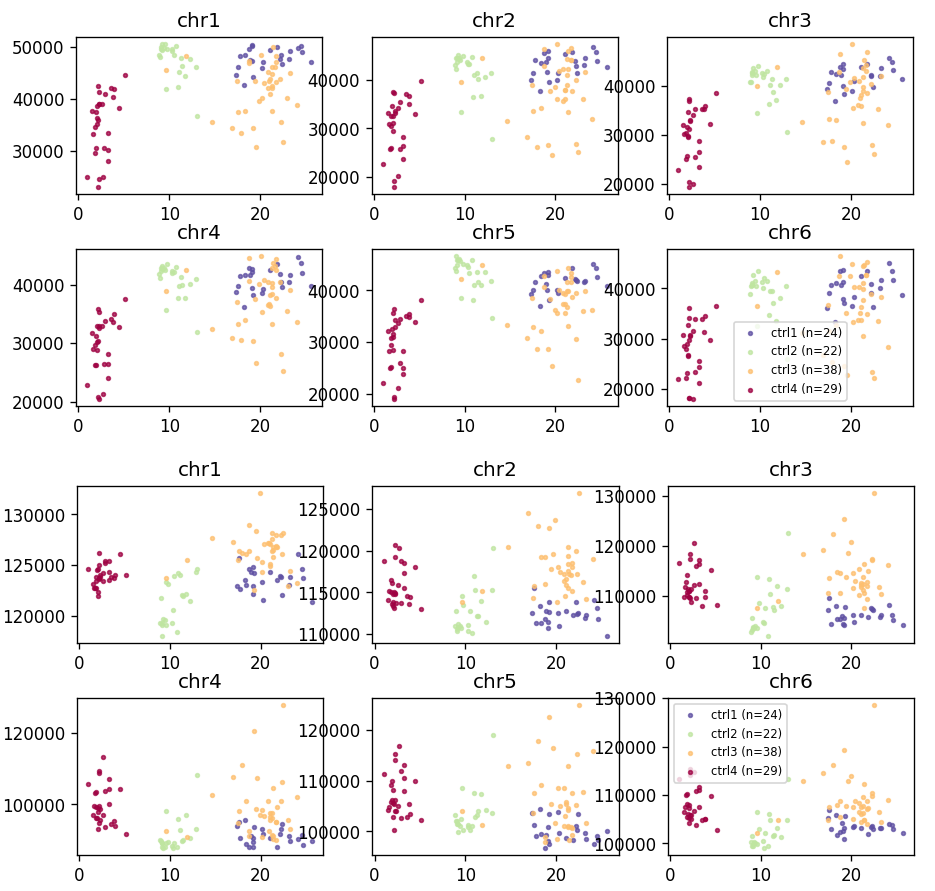

Supplementary Figure 15. Plots of mean sequencing depth (x-axis) versus standard deviation of segment coverage (y-axis) in the four control cfDNA datasets (ctrls1-4), for chromosomes 1-6.
